# Zonal human hepatocytes are differentially permissive to *Plasmodium falciparum* malaria parasites

**DOI:** 10.1101/2020.06.29.175968

**Authors:** Annie S.P. Yang, Youri M. van Waardenburg, Marga van de Vegte-Bolmer, Geert-Jan A. van Gemert, Wouter Graumans, Johannes H.W. de Wilt, Robert W. Sauerwein

## Abstract

*Plasmodium falciparum (Pf)* is a major cause of malaria. The mosquito-borne parasite asymptomatically infects hepatocytes in the liver. The resulting schizonts undergo massive replication to generate blood-infective merozoites. Liver lobules are zonated: hepatocytes in different zones perform differential metabolic functions. In search for specific host conditions that affect infectability, we studied the *Pf* parasite liver stage development in relation to the metabolic heterogeneity of fresh human hepatocytes. We show selective preference of different *Pf* strains for a minority of zone 3 hepatocytes characterized by the particular presence of glutamine synthetase (hGS). Parasite schizont growth is significantly enhanced by hGS uptake early in development, which showcases an import system at this stage of the parasite life-cycle. In conclusion, *Pf* development is strongly determined by the differential metabolic status in hepatocyte subtypes. These findings underscore the importance of detailed understanding of hepatocyte host-*Pf* interactions and may delineate novel pathways for intervention strategies.

## Introduction

Malaria is a devastating mosquito-borne disease responsible for approximately 220 million clinical cases and 430,000 deaths annually [1]. It is caused by the parasites of the *Plasmodium* genus, of which *P. falciparum* (*Pf*) is responsible for most of the disease burden. The infection begins with deposition of sporozoites in the skin by blood-feeding infected mosquitoes. Subsequently low numbers of deposited sporozoites invade, differentiate and massively multiply (as schizont forms) inside hepatocytes followed by release of blood-infective merozoites into the circulation [2]. Cycles of asexual parasite multiplication in circulating red blood cells are responsible for malaria morbidity and mortality [2].

Interaction between host hepatocytes and parasites, especially intracellular host factors that influence parasite development are poorly understood. Studies of *Pf* parasites and host cells, primarily studied with the NF54 strain, are hampered by low parasite infection rates of *in vitro* cultured hepatocytes [3]. Large “omics-based” approaches have been previously applied in rodent malaria models [4, 5], but translational relevance to *Pf* parasites suffers from important biological differences between the two species i.e. the much shorter liver stage of 48 hours (rodent models) as compared to 7 days (*Pf*).

The liver is a complex organ, composed of functional lobules where hepatocytes express distinct metabolic functions related to their zonal location (Z) [6]. Hepatocytes of a particular zone do express specific sets of genes reflective of their differential tasks related to glucose metabolism [7–9]. As such, periportal Z1 is involved in gluconeogenesis during homeostasis while Z3 generates energy through glycolysis [10, 11]. Glucokinase (GK) is an important cytoplasmic enzyme of glycolysis, catalyzing the first step of converting glucose to glucose-6-phosphate [12, 13]. In a gluconeogenic state i.e. Z1, GK are sequestered away from the cytoplasm, being inhibited by its regulatory protein (GKRP) and adopt nuclear localization [14–16]. Contrastingly in Z3, GK translocates from the nucleus to the cytoplasm to participate in glycolysis. Additionally, Z3 hepatocytes are characterized by the exclusive presence of glutamine synthetase (GS), the only so-called stable marker of liver zonation i.e. not affected by the glucose availability of the host [6]. Z2 are located in between Z1 and 3 with apparent limited ability to perform both gluconeogenesis and glycolysis.

The unique intracellular environments present in hepatocytes of different zones may influence the developmental kinetics of obligate intracellular microbes, including *Pf* parasites. *Pf* liver-stages represent an attractive target for vaccine and/or drug development but progress is hindered by limited knowledge of the molecular and cellular events that occur during this stage [2]. While host-*Pf* interactions have been mostly focused on hepatocyte membrane receptors allowing for *Pf* entry [17–20], the intracellular milieu may have major impact on parasite development. Here, we examined whether freshly isolated zonal hepatocytes characterized by hGK and/or hGS expression express differential permissiveness to a panel of clinical *Pf* isolates and explored possible mechanisms involved.

## Results

### Quantification of zonal hepatocytes from freshly isolated primary human hepatocytes (fPHH)

Using specific antibodies against glucokinase (hGK) and glutamine synthetase (hGS) we quantified the proportion of hepatocyte subpopulations fPHH (Supplementary FigS1 for anti-hGS characterization). Z1 was defined by a stronger nuclear hGK signal compared to Z2 with concomitant lack of hGS signals; Z3 was defined by strong cytoplasmic hGS expression combined with an absence of nuclear hGK (Fig 1A). The fPHH monolayers (Fig 1B) from four different human donors were characterized: the vast majority of cells (90%) were Z2 hepatocytes (hGK+/GS−) with the remaining 10% being made up by Z1 (3%; hGK++/GS−) and Z2 (7%; hGK−/GS+).

**Figure 1:**
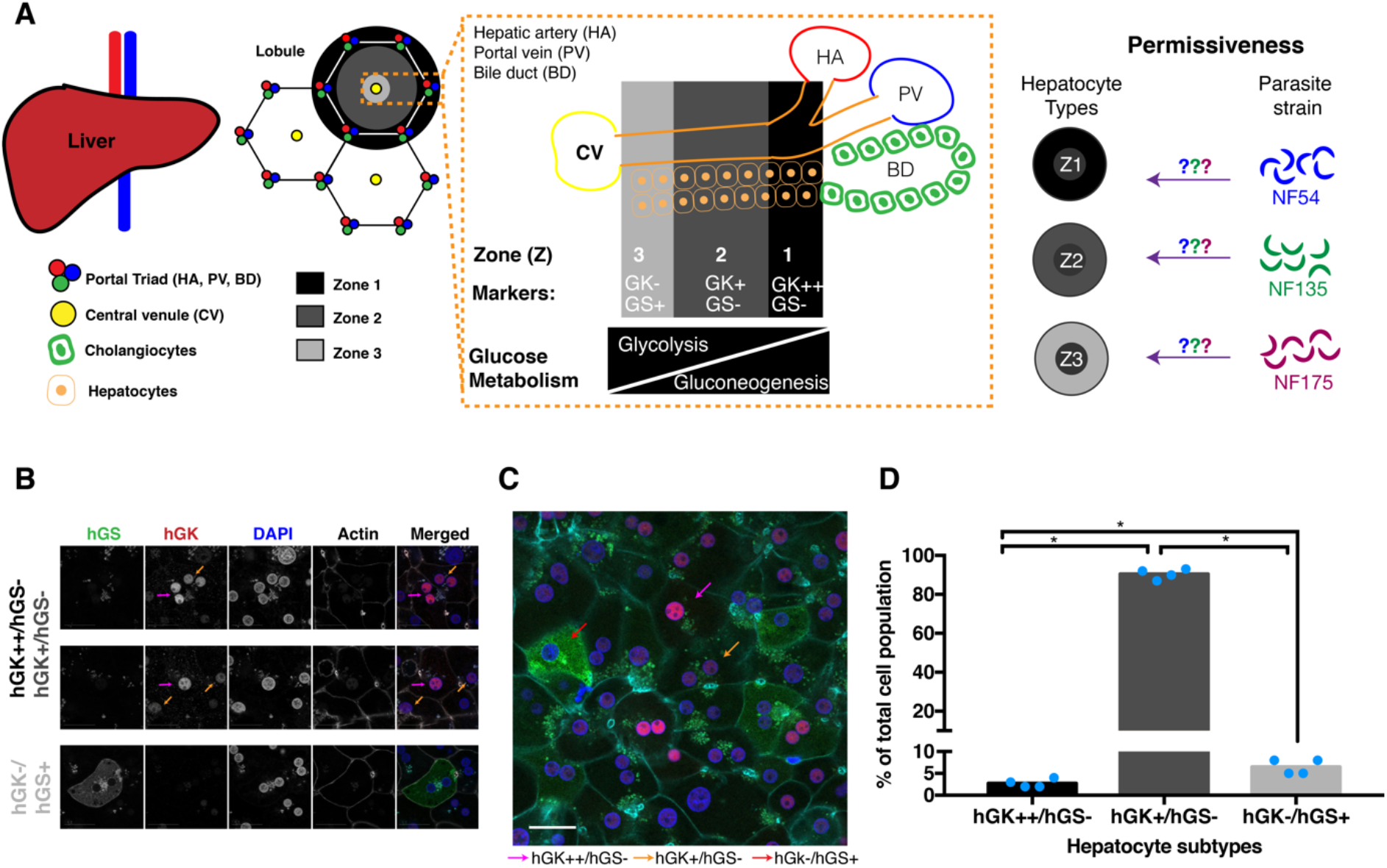
Characterization of zonal hepatocytes from freshly isolated human hepatocytes (fPHH) **A)** Schematic of liver architecture and underlying objective of the study. **B)** Confocal immunofluorescence image of fPHH at 5 days post plating stained with hGK (red), hGS (green), DAPI (blue) and phalloidin (grey). Objective 63x; zoom 2x; scale bar 25 microns. **C)** Confocal immunofluorescence image of a fPHH monolayer at 5 days post plating, stained with hGK (red), hGS (green), DAPI (blue) and phalloidin (cyan). Arrow point towards a typical hepatocyte subpopulation shown by different colours. Objective 40x; scale bar 25 microns. **D)** Percentage of zone 1-3 uninfected hepatocytes day 5 post plating of 4 different donors (blue dots). For each donor, >500 cells were characterized for the final percentages. All different hepatocytes subpopulations are significantly different from each other (Mann-Whitney test: *p = 0.0286*).

### *Permissiveness of hepatocyte subsets for Pf* strains

Established *Pf* strains differed genotypically at a selection of gene loci (Supplementary Fig S2A, B) but did express typical markers used to characterize *Pf-*liver stages (Supplementary Fig S2C). Both NF135 [21, 22] and NF175 [23] showed approximately 4-fold higher numbers of infected fPHH at multiple time points post invasion (p.i.) (Fig 2A) and were significantly larger in size (Fig 2B) compared to NF54.

**Figure 2:**
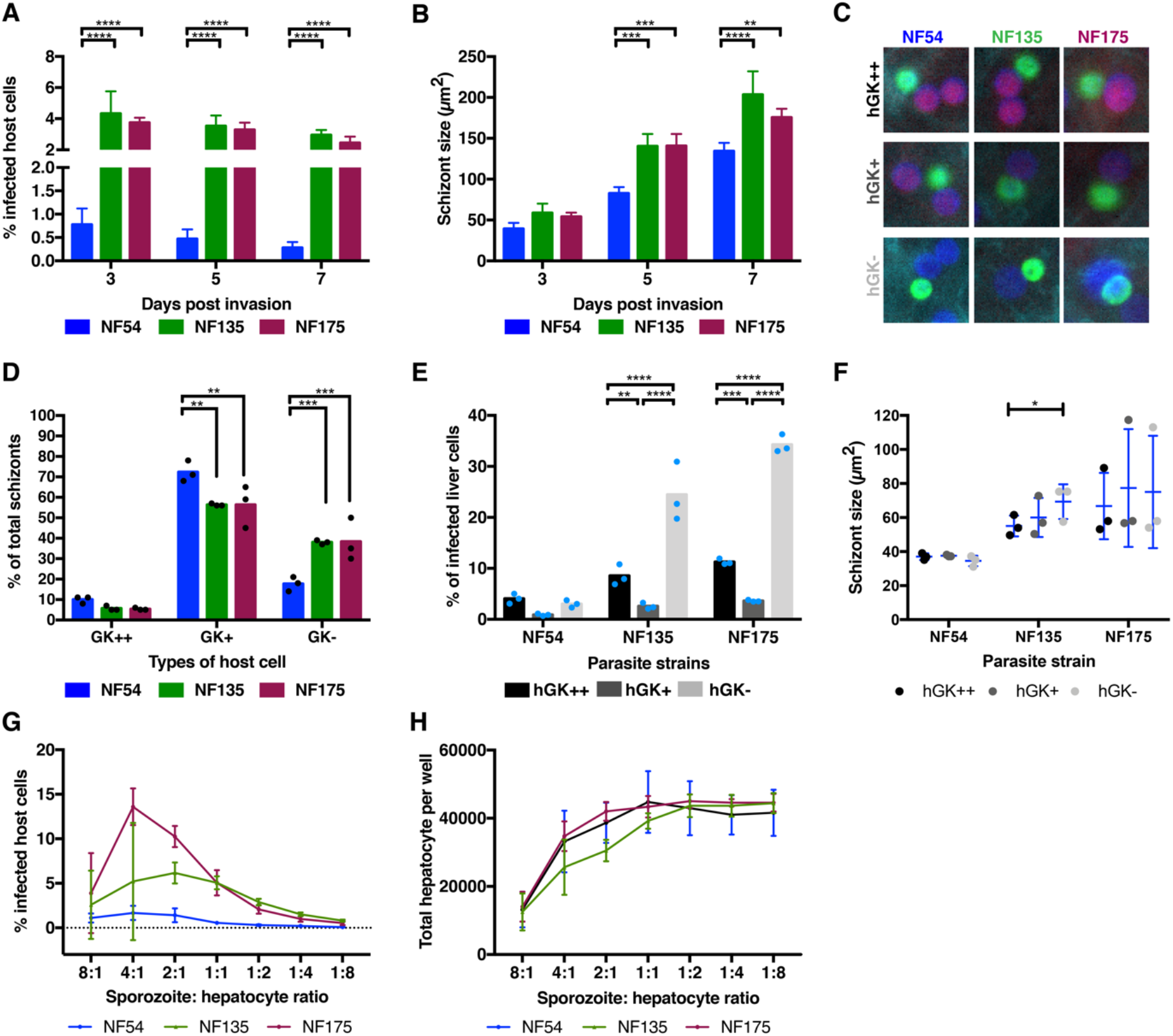
*P. falciparum* development in fPHH and their preference for different hepatocyte subpopulations. **A)** Percentage of hepatocytes with schizonts and **B)** size of schizonts for NF54 (blue), NF135 (green), and NF175 (purple) on days 3, 5 and 7 p.i. with sporozoites to hepatocyte ratio of 1:1. Three biological replicates were performed with two technical replicates in each. At least 100 schizont sizes were measured for each of three biological replicates and the median is plotted above. Tukey’s multiple comparison: ** = 0.0024; *** = 0.0003; and **** = <0.0001. **C)** Representative immunofluorescence images showing the different sub-population of infected fPHH: hGK (red), Dapi (blue) and phalloidin (cyan). Parasites are stained with PfGAPDH (green). Images taken on the Leica High Content at 20x magnification and then cropped in Adobe Photoshop. **D)** Percentage schizonts (from total number of schizonts in a well) located in the different hepatocyte sub-populations by NF54, NF135, NF175 on day 3 p.i.. Each dot represents a biological replicate. For each parasite line, at least 100 schizonts were characterized for the final percentage. Tukey’s multiple comparison test was performed. Percentage of schizonts in hGK++ hepatocytes is not different between the three parasite strains. Percentage of schizonts in hGK+ for NF54 is significantly different from NF135 (*p = 0.0049*) and NF175 (*p = 0.0049*). Percentage of schizonts in hGK- hepatocytes for NF54 is significantly different from NF135 (*p = 0.0006*) and NF175 (*p = 0.0.0005)*. **E)** Percentage of hGK++ (black), hGK+ (dark grey) and hGK- (light grey) hepatocytes that are infected with different *Pf* strains on day 3 p.i.. This is calculated by dividing the number of schizonts by the actual number of hepatocytes in each subpopulation. Each dot represents a different liver donor (same as used in Figure 2D). The percentage of hGK++ (Z1) and hGK- (Z3) hepatocytes infected with parasites were significantly different for NF135 (Dunnett’s multiple comparison: *p = 0.0404* and *p = 0.0001*) and NF175 (Dunnett’s multiple comparison: *p = 0.0015* and *p = 0.0001*) compared to NF54. The percentage of hGK+ (Z2) hepatocytes infected with parasites were not significantly different between the strains. Tukey’s multiple comparison test was used for comparisons within a strain. For NF135, there are significant differences between the percentage of infected hepatocytes within the different zones (hGK++ versus hGK+ *p = 0.0021*, hGK+ versus hGK- *p = <0.0001*, and hGK++ versus hGK- *p = <0.0001*). For NF175, there are also significant differences between the percentage of infected hepatocytes within the different zones (hGK++ versus hGK+ - *p = 0.0003*, hGK+ versus hGK- *p = <0.0001*, and hGK++ versus hGK- *p = <0.0001*). There is no difference in the percentage of infected hepatocytes in the different zones for NF54. **F)** Size of schizonts in hGK++, hGK+ and hGK- hepatocytes on day 3 post invasion for the different *Pf* strains. Tukey’s multiple comparison test was used to show that NF135 hGK++ and hGK- parasites are significantly different (*p = 0.0430*). **G)** Percentage of fPHH with NF54, NF135, and NF175 schizonts on day 5 p.i., under different parasite to hepatocyte ratios (starting from 8 sporozoites to 1 hepatocyte to 1 sporozoites to 8 hepatocytes). Two biological replicates were performed with two technical replicates in each. **H)** Total number of hepatocytes per well for NF54, NF135 and NF175 on day 5 p.i., under different parasite to hepatocyte ratios (starting from 8 sporozoites to 1 hepatocyte to 1 sporozoites to 8 hepatocytes). Two biological replicates were performed with two technical replicates in each.

Next we determined the percentage of *Pf* strains located in different hepatocyte subpopulations on day 3 p.i. (Fig 2C). The percentages of schizonts in hGK++/hGS- (Z1) were similar for all 3 strains. Most schizonts were found in hGK+/hGS- (Z2) cells, representing the majority of cells (Fig 2D). NF135 and NF175 showed relatively higher preference for hGK−/hGS+ (Z3) with up to 35% infection of these cells (Fig 2E). Furthermore, NF135 schizonts present in hGK++ hepatocytes were significantly smaller than those present hGK- (Fig 2F). NF54 showed slight preference for both hGK−/hGS+ (Z3; 3%) and hGK++/hGS- (Z1; 4%) hepatocytes, but overall infection rates and schizont size across the zones were low (Fig 2E, F). The observed distribution of infection may be the reflection of limited availability of preferred fPHHs. Therefore, graded numbers of sporozoites were added to fPHHs and infection rates were determined on day 5 p.i.. Infection rates peaked at sporozoites: hepatocytes ratio of 4:1 for all three strains with higher ratios resulting in a decrease in the number of infected cells, most likely due to deterioration of the monolayer (Fig 2G, H).

### Differential usage of human glutamine synthetase by Pf strains

As the majority of the parasite strains appeared to show a strong preference for hGK−/hGS+ (Z3) cells, we further investigated a possible role of hGS in parasite development because of its exclusive presence in this subset. In uninfected hepatocytes, hGS molecules were uniformly distributed in the cytoplasm but in infected cells, they clustered around the periphery and center of developing NF135 and NF175 schizonts (Fig 3A, B). This condense and centralized accumulation of intra-parasitic hGS further transformed into network-like structures between days 5 to 7. Interestingly, such distribution was not found for NF54 schizonts (Fig 3C).

**Figure 3:**
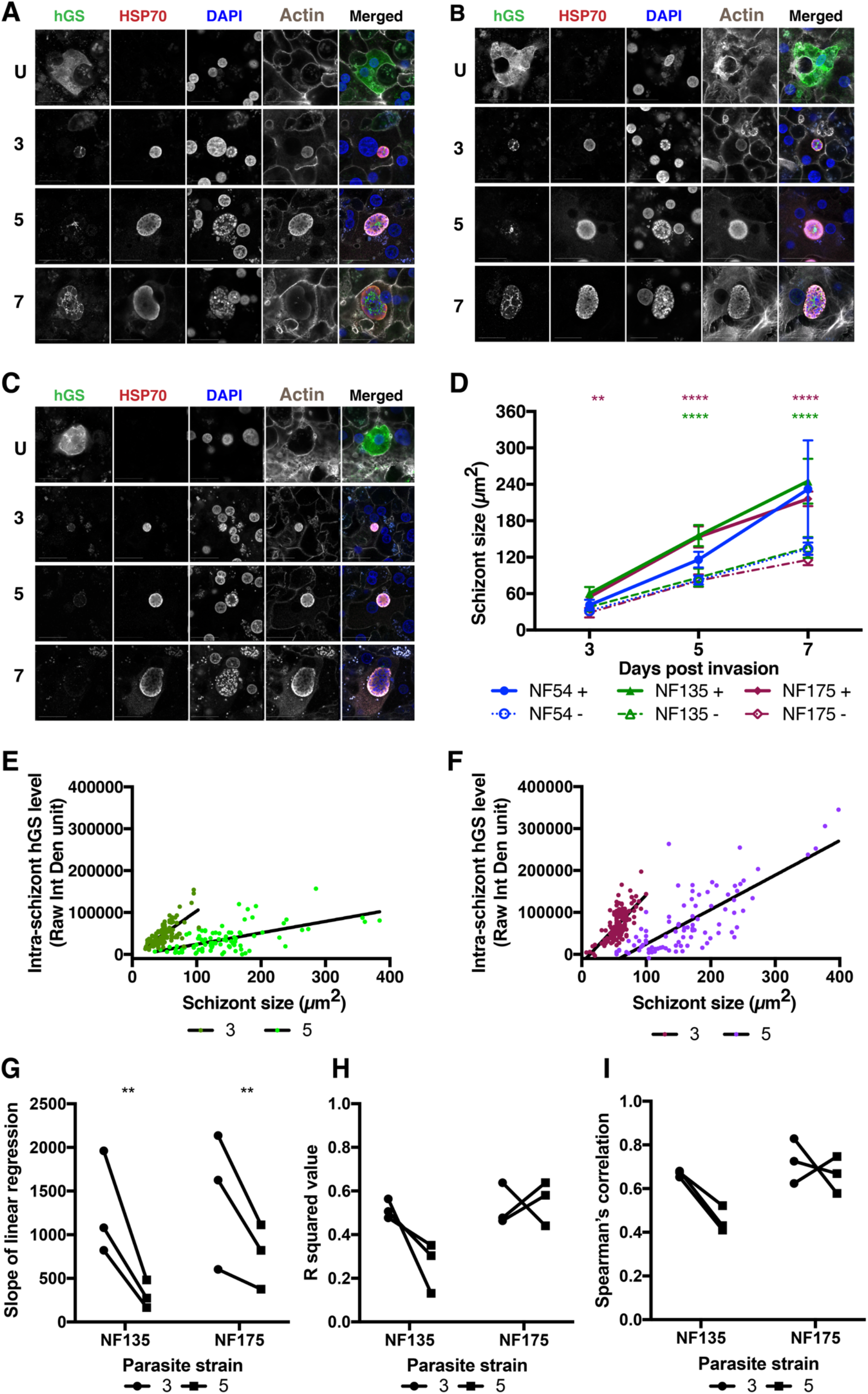
Distribution of human glutamine synthetase (hGS) in different *Pf* strains. **A-C)** hGS interaction with NF135 (A), NF175 (B) and NF54 (C) schizonts on days 3 (top), 5 (middle) and 7 (bottom) p.i.. hGS (green), *Pf*HSP70 (red), DAPI (blue) and phalloidin (grey). The U panel is composed an uninfected zone 3 hepatocyte on day 3 p.i.. Scale bar is 25 microns. **D)** Median size of NF135 (green), NF175 (purple) and NF54 (blue) schizonts, divided based on the presence (solid line) or absence (dashed line) of hGS staining pattern. At least 100 schizonts were measured for each of three biological replicates and the median is plotted above. Sidak’s multiple comparison test was performed (** = p is 0.0052; **** = p is < 0.0001). See Supplementary Fig S4 for raw measurements. **E)** The total hGS signal within intracellular NF135 and **F)**NF175 schizonts on day 3 and 5. Each graph represents a biological replicate where each dot represents an individual schizont. **G)** The slope of the linear regression of the **E)** and **F)** of NF135 and NF175 on day 3 (circle) and day 5 (square) p.i.. **H)** The R squared value of the predicted linear relationship of hGS levels versus schizont size on day 3 and day 5 p.i.. **I)** The Spearman’s correlation coefficient between hGS levels and schizont size of NF135 and NF175 on day 3 and day 5 p.i.. (For G-I: Each dot represents one biological replicate. See Supplementary Figure 3-5 for raw data).

Developing schizonts were categorized by strain for the presence and distinctive staining pattern of hGS (Fig 3D). hGS+ NF135 and NF175 schizonts were significantly larger than their hGS- counterparts at any timepoint p.i.. This could not be studied for NF54 given only very few GS+ schizonts were found from day 5 p.i. onwards. The relationship between intra-schizont hGS levels and size was examined for NF135 and NF175 (Fig 3E, F). Linear regression lines for each strain showed steeper slopes on day 3 than 5 (Fig 3G). Similarly, hGS levels strongly correlated with schizont size (Fig 3H, I) in particular on day 3.

### Effect of GS inhibitors on intra-hepatic parasite development

To confirm the functionality of intra-schizont hGS and its involvement in size regulation, fPHH monolayers infected with either NF135 (hGS benefit) or NF54 (no hGS benefit) were incubated with a panel hGS inhibitors with different modes of action including Glufosinate-ammonium (GA – also known as phosphinothricin) [24, 25], L-methionine sulfoximine (LMS) and 3-aminoimidazole [1,2-a] pyridine (AIP) [26, 27]. LMS and GA compete with the substrate glutamate for the active site; the more recently discovered inhibitor AIP [28, 29] binds to the ATP binding site of the enzyme. AIP significantly reduced the size of developing NF135 schizonts whereas both GA and LMS were inactive (Figure 4A; Supplementary Fig S6, 7). NF54 schizonts were not affected. Importantly, the AIP-induced size reductions were exclusively observed in hGS positive schizonts, strengthening the functionality of hGS for parasite growth (Fig 4B). Both GA and LMS showed no activity (Supplementary Figure S6, 7), which may be due to active metabolization in Z3 hepatocytes [30].

**Figure 4:**
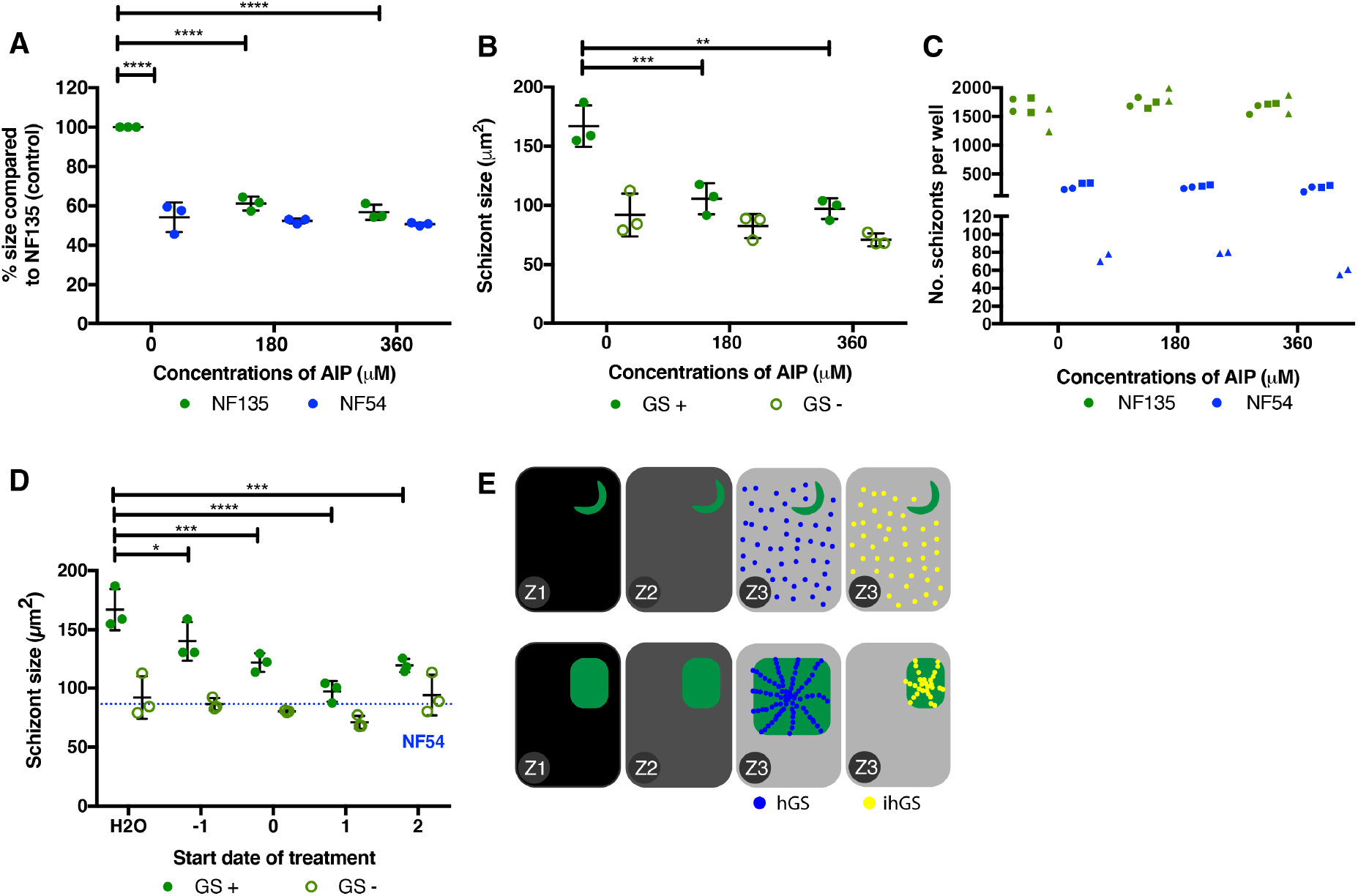
Effect of GS inhibitors on NF54 and NF135. **A)** Normalized percentage of intracellular schizont size of NF54 (blue) and NF135 (green) after treatment of GS inhibitors at different concentrations of AIP. Median size of the schizonts were normalized to that of H2O-treated NF135 (which is set at 100%). AIP was added one day post invasion and kept on for 48 hours. Each dot represents a biological replicate where ≥100 schizonts were measured on day 5 p.i.. For comparisons between NF54 and NF135, Sidak’s multiple comparison test was performed where *** = 0.0005 and ****=<0.0001. Tukey’s multiple comparison test was used for comparisons between conditions within NF135 where * = 0.0135 (LMS) and 0.0401 (GA); and **** = <0.0001. **B)** Size difference between the GS positive (closed circle) and negative (open circle) NF135 schizonts after treatment of AIP. Each dot represents the median of a biological replicate where ≥100 schizonts were measured. Dunnett’s multiple comparison test with *** = 0.0001. **C)** Number of schizonts per well for NF135 (green) and NF54 (blue). Three biological replicates (circle, square and triangle), each with two technical replicates were performed. **D)** The difference in size between GS positive (closed circle) and negative (open circle) NF135 schizonts (day 5 p.i.) after treatment of AIP (360 μM) on different days pre - and post-invasion. Each dot represents the median size of ≥100 schizonts. The GS positive schizont size when treated with AIP were significantly smaller than the water control for day -1 (*p = 0.0348*), day 0 (*p = 0.0005*), day 1 (*p = <0.0001*) and day 2 (*p = 0.0003*) using Tukey’s multiple comparison test. The blue line represents the median schizont size of NF54 when treated with the water control. **I)**Summary of observed effects of hGS on parasite development. Parasites disproportionally prefer Z3 hepatocytes to make use of exclusive presence of hGS. Ability to uptake functional hGS early after invasion results in relatively larger schizonts; uptake of inhibited hGS (ihGS) reduces schizont size. (See Supplementary Figures S7-9 for raw measurements)

Finally, we examined the timing of functional hGS on schizont size. AIP was added for 48 hours to NF135 infected fPHH culture before and after parasite invasion p.i.. There was a steady decrease of intracellular schizont size from day -1 onwards with maximum reduction on day 1 p.i. (Fig 4D). Similarly, size differences between hGS positive and negative schizonts gradually decreased from day -1 onwards (Fig 4D). The relative larger effect of AIP on day 1 p.i. suggests early modulation of schizonts size by hGS; it fits well with the immunofluorescence data regarding localization (Figure 3A) and the strong correlation of hGS positive NF135 and NF175 parasites on day 3 rather than day 5 (Fig 3G-I).

## Discussion

In this study, we show that intra-cellular *Pf* development in the liver is strongly guided by the glucose metabolic status of particular hepatocyte subpopulations. The glycolytic mode and the exclusive presence of hGS in Z3 hepatocytes are beneficial for intra-hepatic stages of particular *Pf* strains as reflected by the larger schizont sizes (NF135 and NF175). The importance of host glucose metabolic status is in agreement with previous findings where *Plasmodium* parasites modulate the host energy sensor, AMP-activated protein kinase (AMPK) to presumably modulate glucose metabolism during liver stage development [31]. Furthermore, the anti-diabetic drug metformin effectively reduces the size of *Pf* liver schizonts. Metformin promotes binding of hGK to GKRP, resulting in a nuclear localization reminiscence of a Z1/2 phenotype [32], shown to result in smaller schizonts in this study.

The observed differences in schizont size of NF135 and NF175 versus NF54 can be explained by the larger number of hGS+ schizonts of the former strains. hGS negative schizonts of all three strains are comparable in size at all time points p.i.. NF54 differs in infectivity, schizont size, host cell preference and hGS utilization from NF135 and NF175. Despite being able to reach Z3 hepatocytes on day 3 pi, NF54 seems unable to make use of host hGS. This is clearly shown in Figure 2D where the sizes of NF54 parasites are similar, irrespective of the host cell type. Additionally, the lack of hGS positive NF54 on day 5 and 7 is actually suggestive for poor survival in the Z3 environment. It remains elusive why NF54 while present in Z3 cells apparently lacks the capacity to mobilize hGS for its own benefit. Furthermore, NF54 parasite development in all hepatocyte subtypes are inferior than NF135 and NF175. Together, it suggests that the observed differences in intracellular parasite numbers between NF54 and NF135/NF175 is due to an inability to make use of hGS that boost schizont development.

Glutamine, as end product of hGS, has been shown to be critical for rapidly proliferating cells; its nitrogen is needed for purines (adenosine and guanine) involved in DNA replication and as building block for newly synthesized proteins and lipids [33, 34]. In this study, this results in the generation of larger schizonts presumably due to more robust DNA, protein and lipid production. The uptake of hGS by certain parasite strains is indicative of an import pathway which may also be involved in the direct transport of other host factors into the parasite. Understanding the timing and mechanism of this pathway may prove to be useful as a direct delivery platform of anti-malarials, bypassing the parasitophorous and parasite membranes. This may be particularly important as *Pf* liver stages represent an attractive target for clinical interventions.

The parasite-induced redistribution of intracellular hGS, may also have direct consequences for hepatocyte survival. Mammalian host cells are equipped with intrinsic detection systems that prevent existence and/or replication of pathogens. Upon detection of the invader, host cells are programmed to create an “anti-microbial defense state” in their cytoplasm [35]. Recently, it has been shown that rodent malaria parasites upregulate cellular inhibitors of apoptosis proteins (cIAPs) in infected hepatocytes [36]. While prevention of cell death by the presence of intracellular parasites has been established in rodent malaria models [37–39], it remains unknown for *Pf* where the development period in hepatocytes takes much longer. High intracellular glutamine concentrations can lead to cell death known as glutamoptosis [40]. It is compelling to speculate that parasites may use hGS both to benefit their own growth and to prevent induction of host death through the accumulation of the glutamine product until their development has been completed.

Schizonts in Z3 grow to larger sizes and would presumably be more likely successfully release larger numbers of infectious progenies into the circulation. It is unknown whether there are zonal differences in maturation, leading to successful release of the parasites into the circulation. Eickel and colleagues previously showed that the majority of the rodent malaria schizonts do not complete their maturation in hepatocytes [41]. Potential zonal differences in successful growth of liver-stages and infection of red blood cells may have implications for therapeutic approaches: there is clear evidence that Z3 hepatocytes are involved in drug metabolism and transport [30, 42]. It would be intriguing to study whether infected Z3 can still maintain its drug metabolic function or whether pre-exposure to anti-malarial drugs has an effect on the growing schizonts and their progenies.

In summary, the combined data show that the developmental kinetics of *Pf* strains is most successful in a minority subset of hGS containing hepatocytes. Future identification of the complete set of host proteins present in hGS positive Z3 hepatocytes will advance understanding of *Pf* interaction with host and may accelerate clinical development of novel intervention strategies against *Pf* liver stages.

## Materials and Methods

### Ethics statement

Primary human liver cells were freshly isolated from remnant surgical material. The samples are anonymized and general approval for use of remnant surgical material was granted in accordance to the Dutch ethical legislation as described in the Medical Research (Human Subjects) Act, and confirmed by the Committee on Research involving Human Subjects, in the region of Arnhem-Nijmegen, the Netherlands

### Antibodies

**Table 1:** Reagents needed for immunofluorescence analysis.

| Antibody | Company | Catalogue no. | Species | Dilution factor |
| --- | --- | --- | --- | --- |
| PfHSP70 | StressMarq<br>Biosciences | SPC-186 | Rabbit | 1:75 |
| PfCSP (2A10) | MR4 | 2A10 | Mouse | 1:500 |
| PfEXP2 | European malaria<br>reagent<br>repository | 7.7 | Mouse | 1:1000 |
| PfGAPDH | European malaria<br>reagent<br>repository | 7.2 | Mouse | 1:50000 |
| PfMSP1 | Sanaria and<br>NIH/NIAID | AD233 | Mouse | 1:100 |
| Human<br>Glucokinase | Abcam | Ab88056 | Rabbit | 1:100 |
| Human<br>Glutamine<br>Synthetase | Abcam | Ab64613 | Mouse | 1:100 |
| Alexa Fluor™<br>647 Phalloidin | Thermofisher<br>Scientific | A22287 | NA | 1:45 |
| Anti-Rabbit<br>Alexa Fluor 594 | Thermofisher<br>Scientific | A11012 | Goat | 1:200 |
| Anti-mouse<br>Alexa Fluor 488 | Thermofisher<br>Scientific | A11029 | Goat | 1:200 |
| DAPI | Thermofisher | D1306 | NA | 1:300 |

### Characterization of genotype of parasite strains

Genomic DNA from different parasite strains were analysed for the presence/absence of GLURP; K1, MAD20, R033 allelic variant of MSP1; and the ICI and FC27 variant of MSP2 as described by McCall et al [22].

### Generation of sporozoites for liver infection

*Pf* asexual and sexual blood stages were cultures in a semi-automatic system as described in [43–45]. *A. stephensi* mosquitoes were reared in the Radboud University Medical Center insectary (Nijmegen, the Netherlands) according to standard operating procedures. Salivary glands from infected mosquitoes were hand dissected and collected in complete William’s B medium (William's E medium with Glutamax [Thermo Fisher, 32551-087], supplemented with 1X insulin/transferrin/selenium [Thermo Fisher, 41400-045], 1mM sodium pyruvate [Thermo Fisher, 11360-070], 1X MEM-NEAA [Thermo Fisher, 11140-035], 2.5 ug/ml Fungizone [Thermo Fisher 15290-018], 200U/ml penicillin/streptomycin [Thermo Fisher 15140-122] and 1.6 μM dexamethasone [Sigma Aldrich D4902-100MG]) without serum. After homogenization using home-made glass grinders, sporozoites were counted in a Burker-Turk chamber using phase contrast microscopy. Immediately before infection of human hepatocytes, the sporozoites are supplemented with heat inactivated human sera (HIHS) at 10% of total volume.

### Primary human hepatocyte infection

Primary human hepatocytes were isolated from patients undergoing elective partial hepatectomy as described by Walk and colleagues [46]. Freshly isolated hepatocytes suspended in complete Williams’ B media were plated in 96 wells at 62.500 cells per well and kept in a 37 degrees Celsius (5% CO2) incubator with daily media refreshments. Two days after plating, dissected sporozoites (day 16-21 post blood meal) were added to the wells at 1:1 ratio in duplicates/triplicates and spun down at 3.000 rpm for 10 minutes on low brakes. Media is refreshed after 3 hours to remove non-invaded sporozoites and then on a daily basis. The sporozoite-infected culture was maintained for 3 or 5 or 7 days after which the cells were fixed with 4% paraformaldehyde (ThermoFisher Scientific: catalogue number 28906) for 10 minutes. The samples were permeabilized using 1% Triton and stained with the various *P. falciparum* or human antibodies listed above.

### GS inhibition

Three GS inhibitors were used to study the effect of GS on parasite infectivity and growth during liver stage development: 3-aminoimidazo[1,2-a]pyridine (Sigma Aldrich; Cat no. 685755; AIP), Glufosinate Ammonium (Sigma Aldrich; Cat no. 45520; GA) and L-Methionine sulfoximine (Sigma Aldrich; Cat no. M5379; LMS). They were tested in two concentrations: AIP (0.18 and 0.36 mM), GA (0.20 and 0.40 mM) and LMS (0.14 and 0.28 mM). Treatment period is for 48 hours with one media refreshment in between after which the culture is returned to complete William’s B media with 10% HIHS until day 5 post invasion where the samples are fixed and stained for analysis by microscopy.

### Microscopy

For this study, the Leica DMI6000B high content microscope was used for tiling 96 wells for determination of infection rate. For each well, a tile size of 9×9 were obtained at 20x objectives. The Zeiss LSM880 with Airyscan at 63x objectives (oil) and 2x zoom were used for detailed images.

### Data analysis using FIJI [47]

#### Infection rate

the tile consisting of 81 smaller images were merged in FIJI and saved as tiff files. Merged tiff files were opened in Adobe Photoshop and manually counted based on PfHSP70 positivity. For NF135 and NF175, only half the final tile (i.e. 40.5 images) were counted and then number of parasites were multiplied by two to get a final total number of parasites per well. Due to the lower infection rate of NF54, the whole tile is counted. Number of hepatic nuclei were counted for 1% of the total image and then multiplied by 100 to get final figure. Infection rate is calculated as total number of parasites (per well) divided by total number of host cells multiplied by 100.

#### Measurement of schizont size

Images obtained on the high content microscope were opened in FIJI. Random images were chosen until 100 parasites were measured. Parasites were selected via the region of interest (ROI) tool using PfHSP70 positivity (red channel) and measured. In the cases, where the hGS intensity is required, the ROIs determined using PfHSP70 are masked onto the hGS channel (green) and then the measured. The RawIntDen values give the total signal measured in the ROI.

#### Quantifying intracellular hGS by immunofluorescence

For each of the image measured in the previous section, a total background intensity in the green channel (as hGS is labelled with Alexa-488) was determined using the region of interest (ROI). A background intensity per area (x) was determined (total intensity of whole image divided by area size of the image). A total background intensity for the parasite was determined by using the formula x multiplied by the measured size of the parasite. This is shown as the background (brown line) in Supplementary Fig S4A, B, D, E, G, H – the larger the parasite, the more total background in theory. Actual green intensity within a parasite is determined by using the ROI tool to just select the parasite in question (the “measured” line in Supplementary Fig S4A, B, D, E, G, H). The final “real” intensity of the parasite is calculated by subtracting the Background value from the Measured value and is what is plotted in Supplementary Fig S5.

### Statistical Analysis

For the majority of the experiments, three biological replicates were performed with either two or three technical replicates (depending on the availability of host cells and parasites). All statistical tests were performed using Prism 7. This includes calculating slope and correlation relationships. See figure legends for details of statistical tests.

## Acknowledgments

We are grateful for R. Stoter, R. Heutink, J. Klaasen, A. Pouwelsen, L. Pelser-Posthumus and J. Kuhnen of the Malaria Unit at the Radboud University Medical Center for parasite, mosquito and sporozoite production. We would like to thank the Microscopic Imaging Center (MIC) of the Radboud University for access to the facilities. Additionally, we would like to thank T. Kooij, N. Proellochs and M. McCall for providing critical feedback on the project as well as R.P. van Rij, K. Dechering and T. Bousema for reviewing the manuscript.

## Author contributions

A.S.P.Y, Y.M.W, G.J.G, M.V, and W.G performed the experiments. J.H.W dW coordinated the collection of fresh human liver segments. A.S.P.Y and Y.M.W collected the data and performed the analysis. A.S.P.Y, Y.M.W and R.W.S were involved in the conceptualization and writing of the manuscript

