## Supplementary figure 1-8 for "Zonal human hepatocytes are differentially permissive to *Plasmodium falciparum* malaria parasites"

**Table of Contents**

**Supplementary Figure 1:** Validation of specificity of monoclonal antibody against human Glutamine Synthetase versus parasite Glutamine Synthetase.

**Supplementary Figure 2:** Supplementary Figure 2: Characterization of NF54, NF135 and NF175.

**Supplementary Figure 3:** Size differences of NF54, NF135 and NF175 schizonts containing GS and without GS.

**Supplementary Figure 4:** Calculating hGS levels

**Supplementary Figure 5:** Relationship between intra-parasitic hGS levels and size of parasites

**Supplementary Figure 6:** Effect of GS inhibitors on parasite size (high concentration)

**Supplementary Figure 7:** Effect of GS inhibitors on parasite size (low concentration)

**Supplementary Figure 8:** Schizont size of NF135 upon treatment with AIP (0.36 mM) on different days pre and post invasion

**A**

```
Human       MTTSASSHLNKGIKQVYMSLPQGEKVQAMYIWIDGTGEGL---RCKT-RTLDSEPKCVEE     56
Pf          ------------MKSVSF-----SNNAELYEYIKDKKNDVEIVACIITNLLGTYFKCFFY     43
                        :*.* :     .:   :* :*... :.:    *   . *.:  **.

Human       ---------LPEWNFDGSSTLQ-SEGSNSDMYLVP--AAMFRDPFRKDPNKLVLCEVFKY     104
Pf          VKEITLNKLESGFSFDASSIKLCSDTEVSDFFIKVDHSTCYLEECDGKNILNIMCDIKRY     103
                        :.**.**    *: . **:::    :: : :    .    ::*:: :*

Human       NRRP----AETNLRHTCKRIMDMVSNQHPWFGMEQEYTLMGTD---------------GH     145
Pf          NGFDYYKCPRTILKKTCEFVKNEGIADKVCIGNELEFFIFDKVNYSLDEYNTYLKVYDRE     163
            *        .* *::**: : :    ::  :* * *: ::..                .

Human       PF------------------GWPSNGFPGPQGPYYCGVGADR-----------AYGR-DI     175
Pf          SFSCKNDLSSIYGNHVVNKVEPHKDHFNNPNNEYLINDDSKKVKKKSGYFTTDPYDTSNI     223
             *                     .: * .*:. *  . .:.:              *.  :*

Human       VEAHYRACL---YAGVKIAGTNAEVMPAQWEFQIGPCEGISMGDHLWVARFILHRVCEDF     232
Pf          I--KLRICRALNDMNINVQRYHHEVSTSQHEISLKYFDALTNADFLLITKQIIKTTVSSF     281
            :  : * *       .:::   : **  :* *:.:   :.:: .*.* ::: *:: . ..*

Human       GVIATFDPKPIPGNWNGAGCHTNFSTKAMREENGLKYIEEAIE-KLSKRHQYHI------     285
Pf          NRTATFMPKPLVND-NGNGLHCNISLWK--NNKNIFYHNDPSTFFLSKESFYFMYGIVKH     338
            .  *** ***: .: ** * * *:*      :::.: * ::      ***.  *.:

Human       ----RAYDPKGGLDNARRLTGFHETSNINDFSAGVANRSASIRIPRTV-GQEKKGYFEDR     340
Pf          AKALQAFCNATMNSYKRLVPGFETCQKL-FY--SFGSRSAVIRLSLINYSNPSEKRIEFR     395
                :*:      .   * : **.  .::  :  ....*** **:      .: .:  :* *

Human       RPSANCDPFSVTEALIRTCLL----NETGDEPFQYKN-----------------------     373
Pf          LPDCANSPHLVMAAIILAGYDGIKSKEQPLVPFESKDNHFYISSIFSKYVQHPENFNILT     455
             *..  .*. *  *:* :       :*    **: *:

Human       ------------------------------------------------------------     373
Pf          HALEGYESLHTINESPEFKNFFKCEEPQGISFSLVESLDALEKDHAFLTVNNIFTEEMIQ     515

Human       ----------------------------     373
Pf          EYIKFKREEIDAYNKYVNAYDYHLYYEC     543
```

**B**

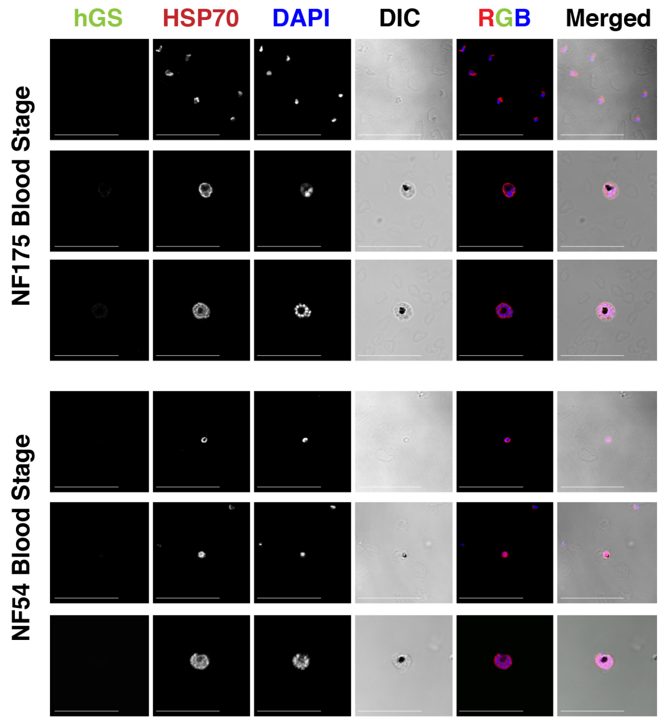


**Supplementary Figure 1: Validation of specificity of monoclonal antibody against human Glutamine Synthetase versus parasite Glutamine Synthetase.**

**A:** Clustal Omega blast of human glutamine synthetase (UniProtKB accession number P15104) and parasite putative glutamine synthetase (PF3D7_0922600). Percentage identity of PfGS compared to hGS is 22.41%. The sequence used to generate the monoclonal antibody is highlighted in yellow and it shows 24.74% identity compared to the aligned region from the parasite GS.

**B:** Top panel: Anti-human glutamine synthetase antibodies (1:100 dilution) on blood stage cultures of NF175 (top panel) and NF54 (bottom panel) respectively showing no reactivity with parasite GS in rings (top), trophozoites (middle) and schizonts (bottom) using the same settings on the Zeiss Airyscan confocal microscope. Scale bar is 25 microns.

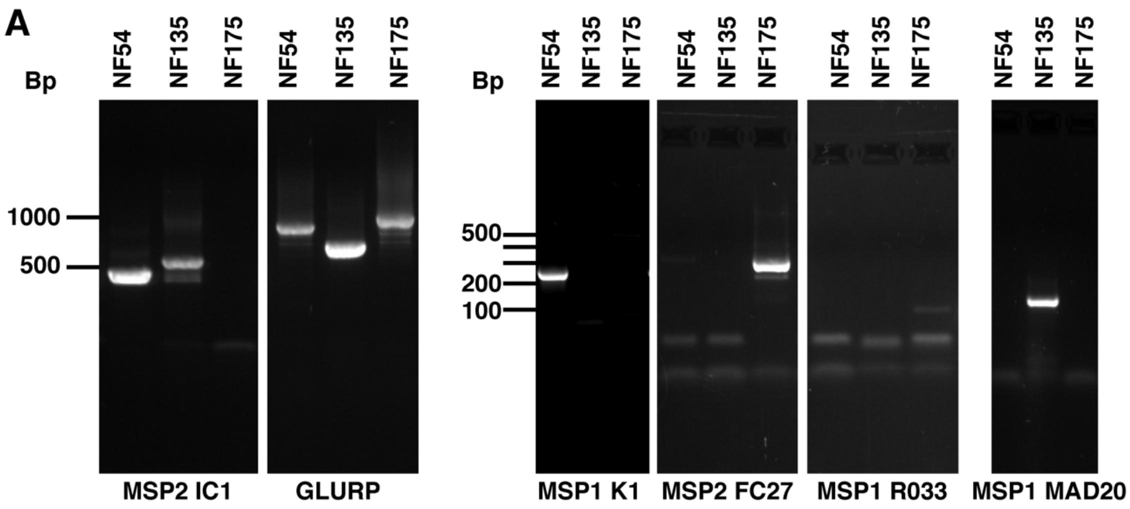


**B**

| Strain | MSP2 IC1 | GLURP | MSP1 K1 | MSP2 FC27 | MSP1 R033 | MSP1 MAD20 |
|---|---|---|---|---|---|---|
| NF54 | 480 | 950 | 220 | neg | neg | neg |
| NF135 | 580 | 680 | neg | neg | neg | 180 |
| NF175 | neg | 990-1000 | neg | 350 | 170 | neg |

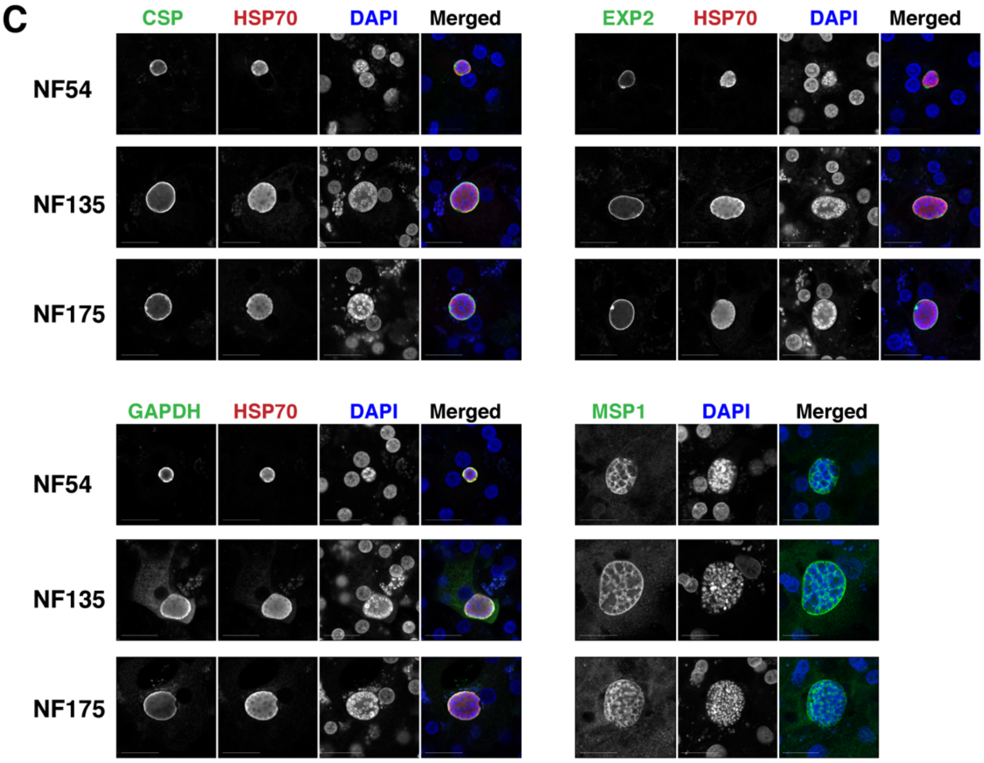


**Supplementary Figure 2: Characterization of NF54, NF135 and NF175**

**A:** Molecular characterization by PCR using established markers: MSP2 IC1, GLURP, MSP1 K1, MSP2 FC27, MSP1 R033, and MSP1 MAD20.

**B:** Markers as in A as well as the sizes for the three parasite strains.

**C:** Expression of typical liver stage protein markers: Circumsporozoite protein (CSP) top left; Exported Protein 2 (EXP2) top right; Glyceraldehyde 3-phosphate dehydrogenase (GAPDH) bottom left; and Merozoite surface protein 1 (MSP1) bottom right. Scale bar is 25 microns.

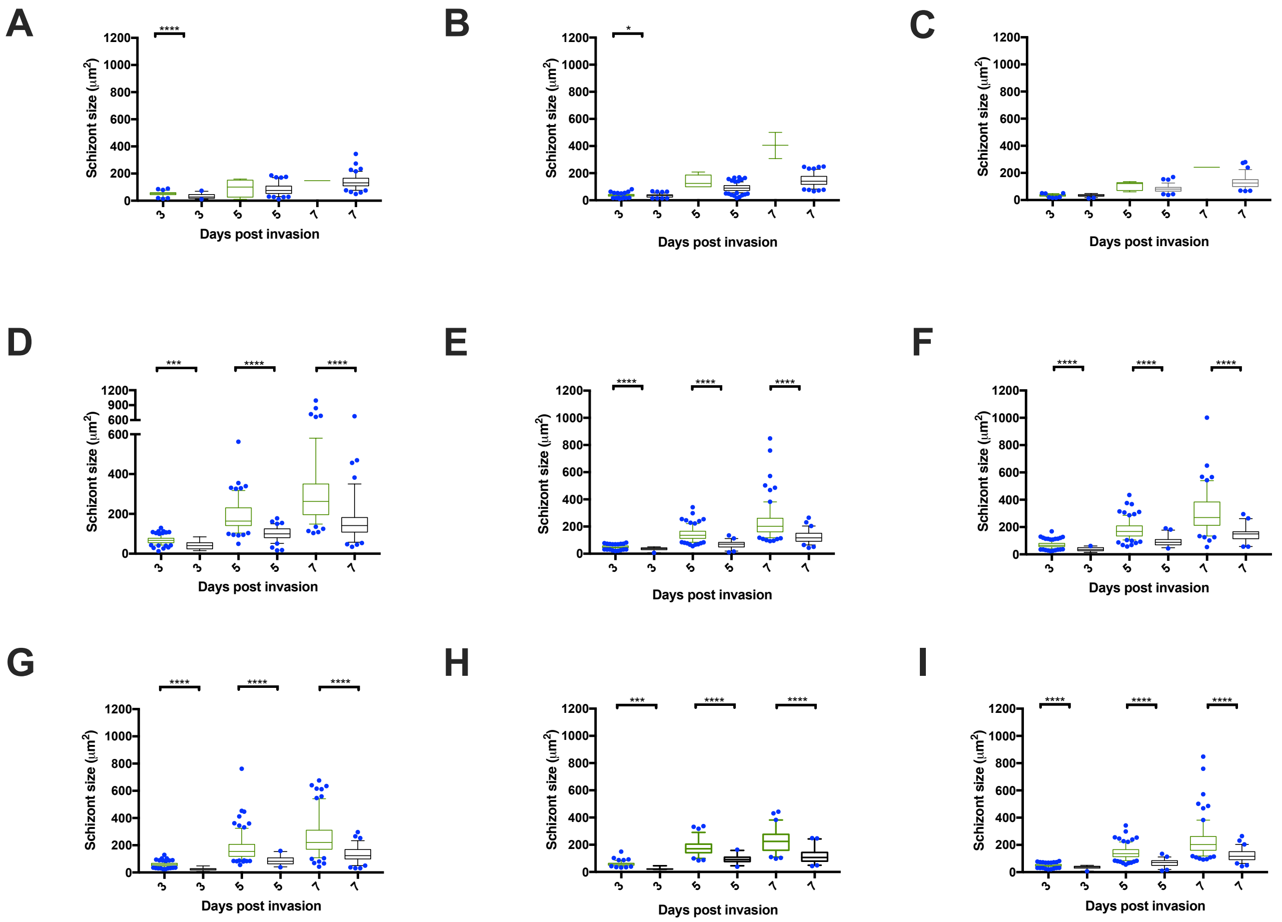


**Supplementary Figure 3: Size differences of NF54, NF135 and NF175 schizonts containing GS (green) and without GS (black).**

**A-C:** Sizes of NF54 schizonts in three different hepatocyte donors. Each graph is a biological replicate where at least 100 schizonts were measured and divided into hGS positive (green border) and negative (black border) groups. Horizontal line within the box plot represents median and the blue dots shows value outside the 5-95% confidence interval. Statistical test used is Mann-Whitney test (also applies for D to I).

**D-F:** Sizes of NF135 schizonts in three different hepatocyte donors. Each graph is a biological replicate where at least 100 schizonts were measured and divided into hGS positive (green border) and negative (black border) groups. Horizontal line within the box plot represents median and the blue dots shows value outside the 5-95% confidence interval.

**G-I:** Sizes of NF175 schizonts in three different hepatocyte donors. Each graph is a biological replicate where at least 100 schizonts were measured and divided into hGS positive (green border) and negative (black border) groups. Horizontal line within the box plot represents median and the blue dots shows value outside the 5-95% confidence interval.

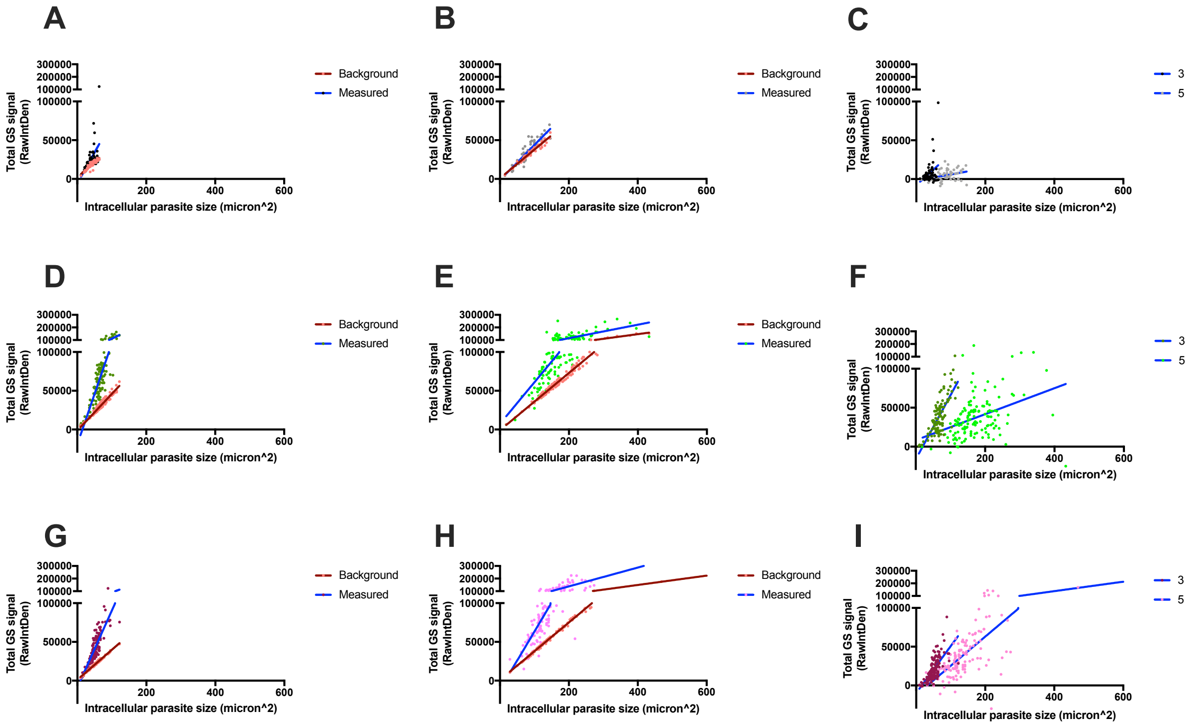


**Supplementary Figure 4: Calculating hGS levels**
In this assay, glutamine synthetase is visualized in the green channel which also has non-specific signal due to the intrinsic autofluorescence feature of the fPHH. Therefore, we calculated a signal per pixel$^2$ value caused by the background green signal: the RawIntDen value divided by the area of the image (pixel$^2$). From there, a predicted background total green signal per parasite was calculated: size of parasite (pixel$^2$) multiplied the background signal (signal per pixel$^2$). This is shown by the orange dots and brown line: the bigger schizonts are, the more total background signal there is. Using the region of interest function of FIJI, we compared this background value to the actual measured green signal in schizonts of NF54 (black for day 3 and grey for day 5), NF135 (dark green for day 3 and light green for day 5) and NF175 (purple for day 3 and pink for day 5). For NF54, the measured dots (black/grey) overlaps with those of the background suggesting that the hGS levels in these parasites are due to background green signal. However, for NF135 and NF175, the measured dots are significantly higher than the predicted background signal, indicating that the signal is due the presence of hGS rather than non-specific background.

For each parasite line, day 3 and 5 schizont sizes were measured (A, B for NF54; D, E for NF135; and G,H for NF175). The final hGS level (measured value minus the background value) are plotted in C, F and I for the different strains.

The background lines for NF54 on day 3 and 5 are not significantly different from the measured line (0.0168 and 0.0114) whereas it is significantly different for NF135 and NF175 (<0.0001).

C: The linear regression line for NF54 on day 3 and day 5 has $R^2$ values of 0.0981 and 0.111 and a Spearman correlation R of 0.096 and 0.3423 respectively.

F: The linear regression line for NF135 on day 3 and day 5 has $R^2$ values of 0.5636 and 0.1312 and a Spearman correlation R of 0.6807 and 0.4306 respectively.

I: The linear regression line for NF175 on day 3 and day 5 has $R^2$ values of 0.4732 and 0.5801 and a Spearman correlation R of 0.315 and 0.6685 respectively.

.

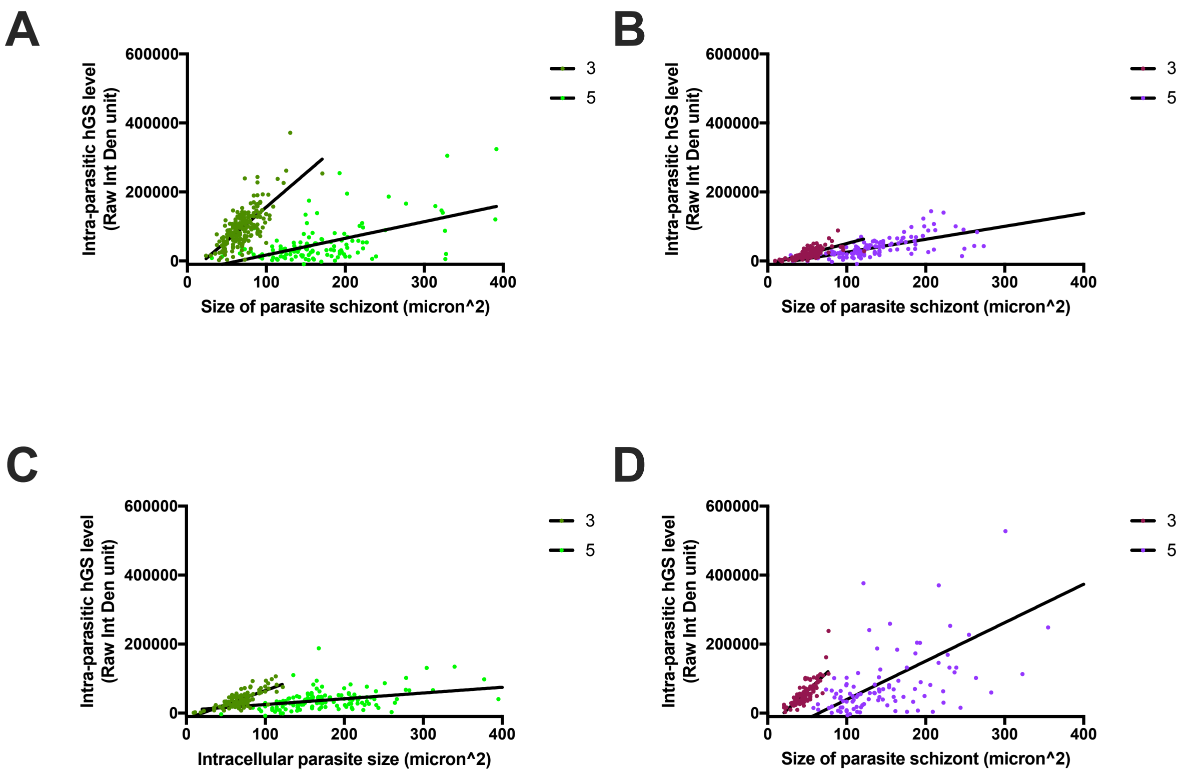


**Supplementary Figure 5: Relationship between intra-parasitic hGS levels and size of schizonts**

A and C: The total GS signal within intracellular NF135 parasite on day 3 and 5. Each graph represents a biological replicate with each dot representing the measurements of one intracellular parasite.

B and D: The total GS signal within intracellular NF175 parasite on day 3 and 5. Each graph represents a biological replicate with each dot representing the measurements of one intracellular parasite.

*The third biological replicate is in main figure 3E and F.*

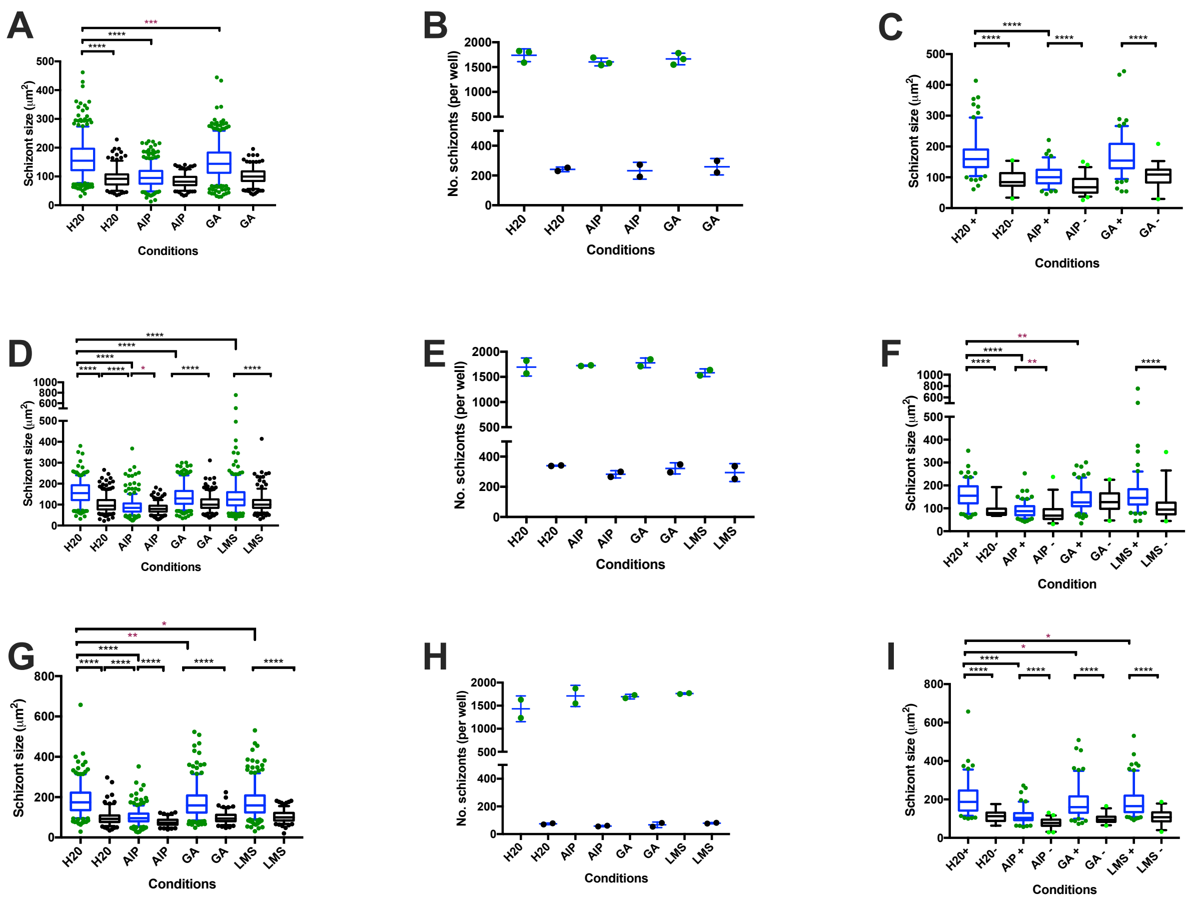


**Supplementary Figure 6: Effect of GS inhibitors on schizont size (higher concentration)**

**A:** Size of intracellular schizont of treated NF54 (black dots and borders) and NF135 (green dots and blue borders) for water control; AIP (360 µM); GA (400 µM) and LMS (280 µM). At least 100 schizonts were measured per conditions. Horizontal line within the box plot represents median and the dots shows value outside the 5-95% confidence interval. Statistical test used is Mann-Whitney test.

**B:** Number of intracellular schizonts of treated NF54 and NF135 per well (technical replicate).

**C:** Size of intracellular schizont of hGS positive (dark green and blue border) and hGS negative (light green and black border) for NF135 treated parasites. At least 100 schizonts were measured per condition and divided into hGS positive and negative groups. Horizontal line within the box plot represents median and the dots shows value outside the 5-95% confidence interval. Statistical test used is Mann-Whitney test

**D:** Same as A but for Biological Replicate 2.

**E:** Same as B but for Biological Replicate 2.

**F:** Same as C but for Biological Replicate 2.

**G:** Same as A but for Biological Replicate 3.

**H:** Same as B but for Biological Replicate 3.

**I:** Same as C but for Biological Replicate 3.

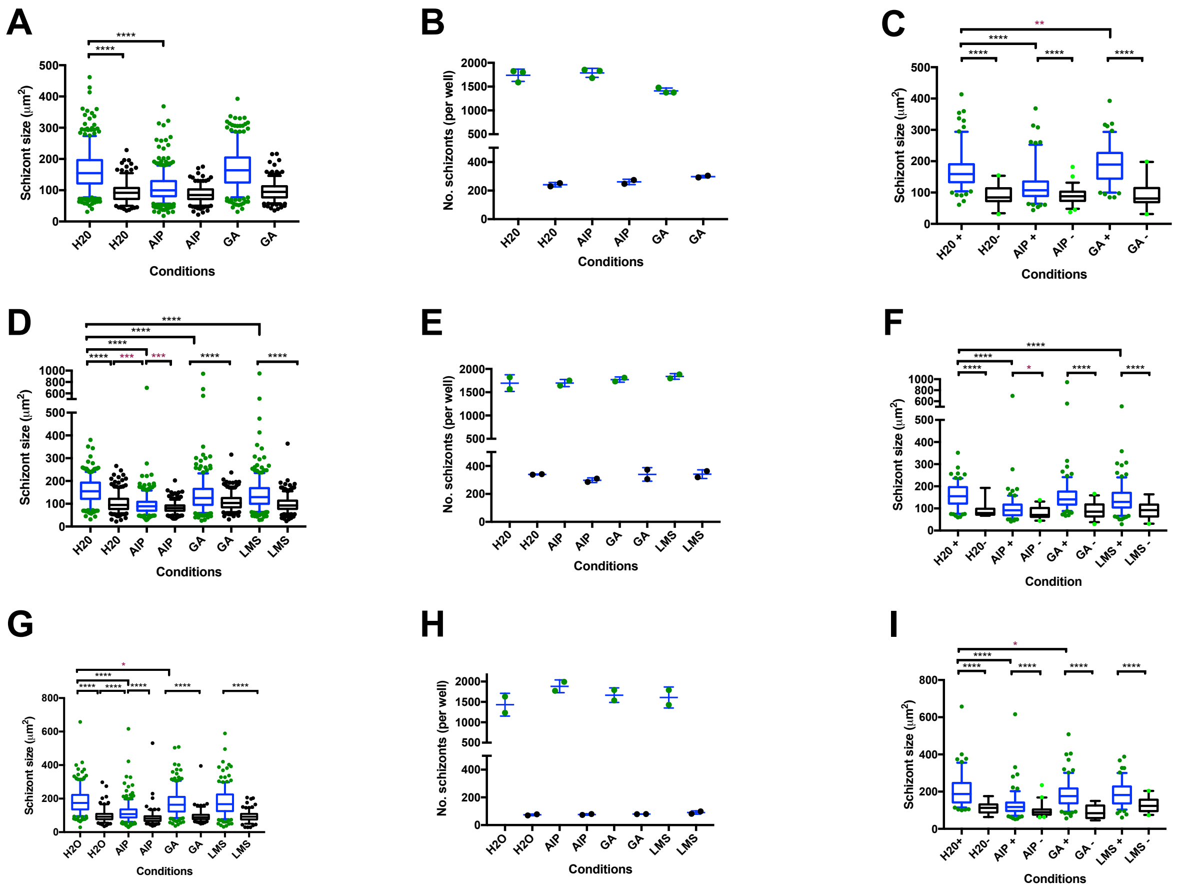

**Supplementary Figure 7: Effect of GS inhibitors on schizont size (lower concentration)**

**A:** Size of intracellular schizont of treated NF54 (black dots and borders) and NF135 (green dots and blue borders) for water control; AIP (108 µM); GA (200 µM) and LMS (140 µM). At least 100 schizonts were measured per condition. Horizontal line within the box plot represents median and the dots shows value outside the 5-95% confidence interval. Statistical test used is Mann-Whitney test.

**B:** Number of intracellular schizonts of treated NF54 and NF135 per well (technical replicate).

**C:** Size of intracellular schizont of hGS positive (dark green and blue border) and hGS negative (light green and black border) for NF135 treated parasites. At least 100 schizonts were measured per condition and divided into hGS positive and negative groups. Horizontal line within the box plot represents median and the dots shows value outside the 5-95% confidence interval. Statistical test used is Mann-Whitney test

**D:** Same as A but for Biological Replicate 2.

**E:** Same as B but for Biological Replicate 2.

**F:** Same as C but for Biological Replicate 2.

**G:** Same as A but for Biological Replicate 3.

**H:** Same as B but for Biological Replicate 3.

**I:** Same as C but for Biological Replicate 3.

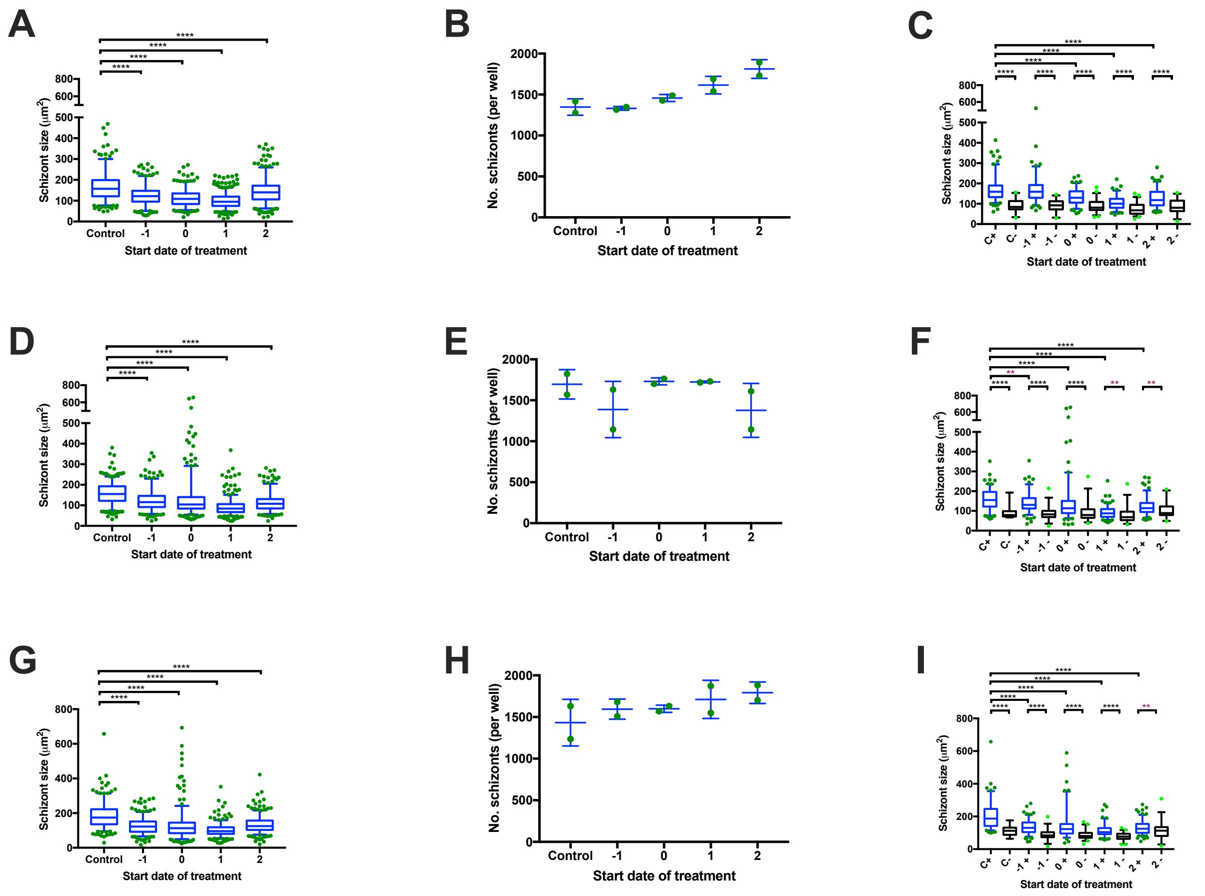


**Supplementary Figure 8. Schizont size of NF135 upon treatment with AIP (360 μM) on different days pre and post invasion**

**A:** Intracellular parasite sizes of NF135 (green) where at least 100 parasites were measured for each condition. Horizontal line within the box plot represents median and the dots shows value outside the 5-95% confidence interval. Mann-Whitney tests were used for the statistical analysis.

**B:** Number of intracellular schizonts of treated NF135 per well (technical replicate).

**C:** Size of intracellular schizont of hGS positive (dark green and blue border) and hGS negative (light green and black border) for NF135 treated parasites. At least 100 schizonts were measured per condition and divided into hGS positive and negative groups. The water control is denoted by the letter C. Horizontal line within the box plot represents median and the dots shows value outside the 5-95% confidence interval. Mann-Whitney tests were used for the statistical analysis.

**D:** Same as A but for Biological Replicate 2.

**E:** Same as B but for Biological Replicate 2.

**F:** Same as C but for Biological Replicate 2.

**G:** Same as A but for Biological Replicate 3.

**H:** Same as B but for Biological Replicate 3.

**I:** Same as C but for Biological Replicate 3.
